# Type IV collagen is essential for proper function of integrin-mediated adhesion in *Drosophila* muscle fibers

**DOI:** 10.1101/318337

**Authors:** András A. Kiss, Nikoletta Popovics, Kiss Márton, Zsolt Boldogkői, Katalin Csiszár, Mátyás Mink

## Abstract

Congenital muscular dystrophy (CMD), a subgroup of myopathies and a genetically and clinically heterogeneous group of inherited muscle disorders is characterized by progressive muscle weakness, fiber size variability, fibrosis, clustered necrotic fibers, and central myonuclei present in regenerating muscle. Type IV collagen (*COL4A1*) mutations have recently been identified in patients with intracerebral, vascular, renal, ophthalmologic pathologies and congenital muscular dystrophy, consistent with diagnoses of Walker–Warburg Syndrome or Muscle–Eye–Brain disease. Morphological characteristics of muscular dystrophy have also been demonstrated *Col4a1* mutant mice. Yet, several aspects of the pathomechanism of COL4A1-associated muscle defects remained largely uncharacterized. Based on the results of genetic, histological, molecular, and biochemical analyses in an allelic series of *Drosophila col4a1* mutants, we provide evidence that *col4a1* mutations associate with severely compromised muscle fibers within the single-layer striated muscle of the common oviduct, characterized by loss of sarcomere structure, disintegration and streaming of Z-discs, and aberrant integrin expression within the M-discs, indicating an essential role for the COL4A1 protein. Features of altered cytoskeletal phenotype include actin bundles traversing over sarcomere units, amorphous actin aggregates, atrophy and aberrant fiber size. The mutant COL4A1-associated defects appear to recapitulate integrin-mediated adhesion phenotypes observed in *Drosophila* by RNA-inhibition. Our results provide insight into the mechanistic details of COL4A1-associated muscle disorders and suggest a role for integrin-collagen interaction in the maintenance of sarcomeres.

## INTRODUCTION

Basement membranes (BMs) are 80-100 nm thick, sheet-like extracellular matrices underlying epithelial and endothelial cells in muscular, neural, vascular and adipose tissues. BMs contain heterogenous composition of major and minor proteins, including type IV collagen, laminin, nidogen/entactin, perlecan, and integrins anchore the BM to the cytoskeleton (Pozzi et al., 2017). Integrity of the BM is a prerequisite for skeletal muscle stability. Positional cloning in humans and targeted gene inactivation in mice revealed that several muscular dystrophy types may develop as the result of the loss of cell-BM anchorage (Sanes 2003). Causative gene mutations were reported in the *laminin-A2* (*LAMA2*) gene, causing LAMA2 (merosin) deficiency (Guicheney et al., 1997; Durbeej and Campbell, 2002). Mutations in the collagen VI genes (*COL6A1*, *COL6A2* and *COL6A3*) were linked to Ullrich CMD, to the milder Bethlem myopathy and to autosomal recessive myosclerosis myopathy (Lampe and Bushby, 2005), while integrin A7 (*ITGA7*) and A5 (*ITGA5*) mutations were shown to be associated to a rare form of CMD (Mayer et al., 1997; Bertini et al., 2011; Hynes 2002).

Mammals harbor three pairs of head-to head oriented type IV collagen genes, *COL4A1* through *COL4A6*, whereas *Drosophila* has one pair, the *col4a1* and *col4a2* genes, in the same genomic organization (Kelemen-Valkony et al., 2012). Expression of the human *COL4A3*, *A4*, *A5* and *A6* genes are restricted both spatially and temporally and confined to the retina, cochlea and kidney. Mutations in the *COL4A3*, *A4* and *A5* genes associate with Alport Syndrome ( Alport 1927; Hudson et al., 2003). Deletions within the *COL4A5* and *COL4A6* genes are also reported to cause diffuse leiomyomatosis, a benign form of tumor-like hypertrophy of the visceral muscle, affecting the oesophagus, the tracheo-bronchial tree, or the female genital tract (Zhou 1993).

Heterotrimers consisting of two COL4A1 and one COL4A2 chains of type IV collagen constitute stochiometrically the most abundant components of nearly all mammalian basement membranes. Mice heterozygous for either missense or exon-skipping mutations of *Col4a1* or *Col4a2*, develop complex, systemic and pleiotropic pathological phenotypes affecting the central nervous, ocular, renal, pulmonary, vascular, reproductive and muscular systems (Gould et al., 2005; van Agtmael et al., 2005; Favor et al., 2007). *COL4A1* or *COL4A2* mutations in humans cause similar cerebral, cerebrovascular, ocular, renal, and muscular pathologies (Kuo et al., 2012). Severe muscular phenotypes were reported in patients with certain *COL4A1* mutations as part of a multi–system disorder referred to as hereditary angiopathy with nephropathy, aneurysms, and muscle cramps (HANAC) with subjects also having elevated serum creatine kinase concentrations (Plaisier et al., 2007; Alamowitch et al., 2007). Some patients with *COL4A1* mutations were also diagnosed with Walker–Warburg Syndrome or Muscle–Eye–Brain disease, a distinct form of CMD (Labelle-Dumais et al., 2011). *Col4a1^G498V/G498V^* homozygous mice are severely affected by muscular dystrophy including muscle mass decrease, fiber atrophy, centronuclear fibers, fibrosis, focal perivascular inflammation and intramuscular hemorrhages (Guiraud 2017).

In HANAC Syndrome, mutations proved to affect multiple putative integrin binding sites within the COL4A1 protein (Parkin et al., 2011; Plaisier et al., 2010). Proper integrin concentration/function was shown to be required for maintenance of the sarcomere structure (Rui et al., 2010). The pivotal role for integrins in myofibril striation was demonstrated by the complete loss of sarcomeres in *Drosophila* integrin null mutants (Volk et al., 1990). In *Drosophila*, the ubiquitous integrin dimer is composed of one of the alpha PS (position-specific) subunits combined with the beta PS protein (Volk et al., 2002). Conditional RNAi knockdown of genes involved in integrin-mediated adhesion, including *talin*, *alpha-actinin*, *integrin-linked kinase*, *alpha PS2* and *beta PS* integrins, revealed a spectrum of phenotypes affecting Z-disc proteins that were dislocated and deposited across the sarcomere, and Z-disc streaming characteristic of myopathic/dystrophic conditions (Perkins et al., 2010).

We have identified an allelic series of conditional, temperature-sensitive *col4a1* mutations in *Drosophila*. The *col4a1^-/-^* homozygotes are embryonic lethal while *col4a1^+/-^* heterozygotes are viable and fertile at permissive temperature of 20°C, but perish at restrictive condition of 29°C. In these mutants, we have demonstrated severe myopathy (Kelemen-Valkony et al., 2012), irregular and thickened BM, detachment of the gut epithelial and visceral muscle cells from the BM (Kelemen-Valkony et al., 2012A), intestinal dysfunction, overexpression of antimicrobial peptides, excess synthesis of hydrogen peroxide and peroxynitrite (Kiss M et al., 2016), furthermore, in epithelial cells of Malpighian tubules, the *Drosophila* secretory organ, fused mitochondria, membrane peroxidation (Kiss AA et al., 2018), actin stress fibers and irregular integrin expression (Kiss AA et al., 2016). Our results indicated that muscular dystrophy may also be present in *col4a1* mutant *Drosophila* (Kiss M et al., 2012).

In order to characterize muscle phenotype in the *col4a1* allelic mutant series, we have determined the mutation sites, in immunohistochemistry experiments focused on the striated oviduct muscle, in the mutant lines we noted aberrant sarcomeres, altered integrin expression and localization, Z-disc disorganization and streaming, fiber size disproportion and atrophy. Results collectively indicate that in mutants, the dystrophic muscle phenotype appears to originate from compromised integrin interactions with aberrant COL4A1, and supports a role for type IV collagen as part of integrin-mediated muscle cell adhesion.

## RESULTS

### Characterization of *col4a1* mutation sites

We have analyzed the DNA sequence of PCR products of the *col4a1* gene using genomic DNA isolated from our series of *col4a1^+/-^* heterozygotes. Consistent with the ethyl-methane-sulfonate (EMS) mutagenesis used to generate these mutants (Kelemen-Valkony et al., 2012), by which the product, *O*-6- ethylguanosine, mispairs with T in the next round of replication, causing a G/C to A/T transition, we identified transition in all mutant loci (Supplementary Fig. S1). Resulting from these transitions glycine substitutions by aspartic acid, glutamic acid or serine were identified in the mutants; hereafter we refer to the mutations as displayed in Table 1. In our prior study we reported five mutations in the *col4a1* gene; beyond the G552D lesion the A1081T amino acid substitution arose by transition, similarly to the G to A transition within the 3’ UTR region of the gene. The K1125N, G1198A amino acid substitutions are results of transversions (Kelemen-Valkony et al., 2012). We recorded these mutation sites in all lines sequenced, therefore concluded that these are carried by the balancer chromosome, *CyRoi*. These balancer chromosome mutations do not contribute to the phenotype, given the most robust phenotypical feature of the mutants, the dominant temperature sensitivity is not influenced by exchange of the *CyRoi* chromosome into the chromosome carrying the *col4a2::GFP* transgene (Kelemen-Valkony et al., 2012).

**Table 1.**
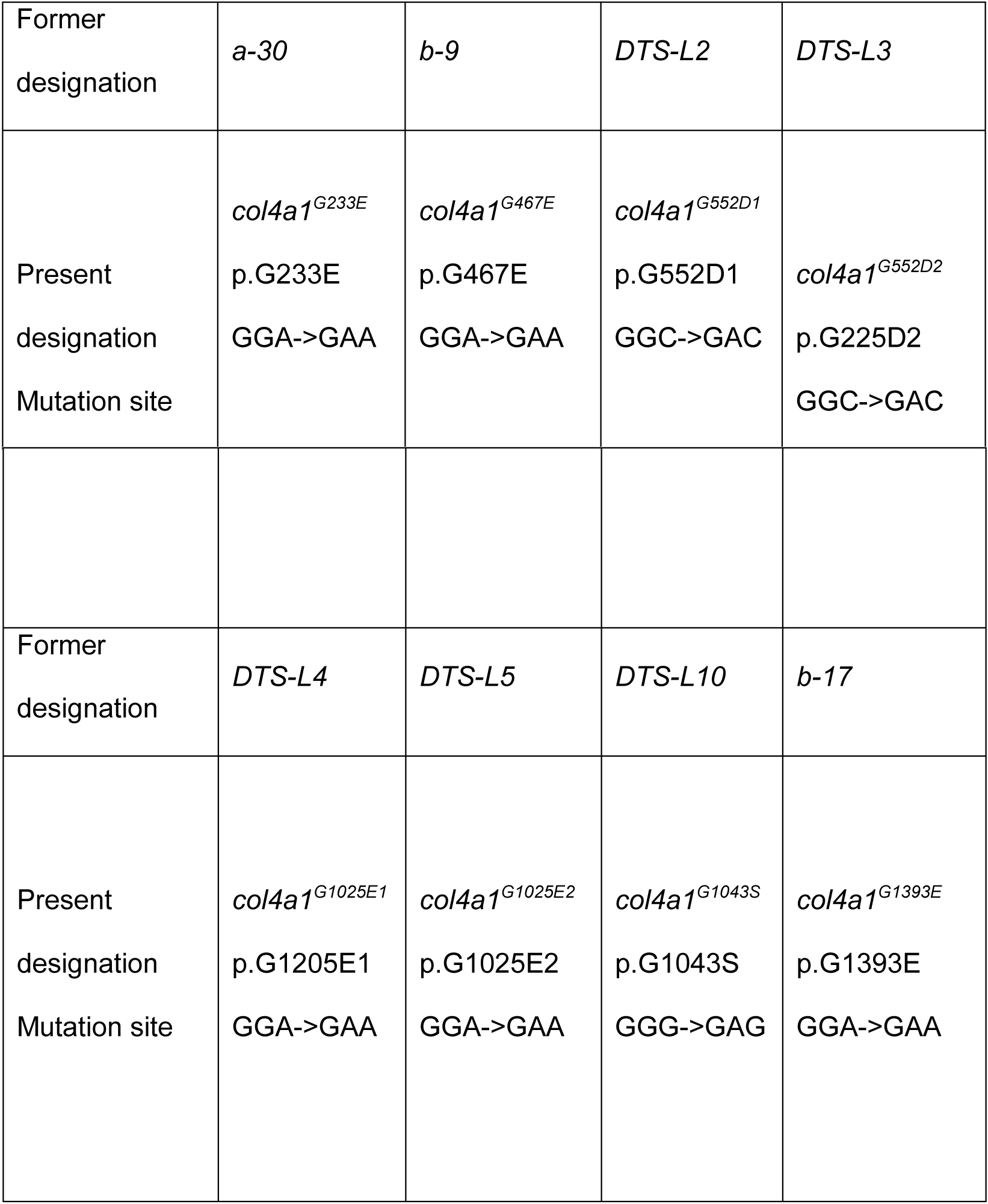
The mutation sites, former and present designation of the mutant loci.

Two pairs of alleles were found to carry the same mutation. The *DTS-L2* and *DTS-L3* lines harbor the same G552D substitution and we refer to these as *col4a1^G552D1^* and *col4a1^G552D2^*. Similarly, the *DTS-L5* and *DTS-L10* lines both carry the G1025E substitution and were designated as *col4a1^G1025E1^* and *col4a1^G1025E2^* alleles. These data confirmed our previous genetic results, for example: The *col4a1^G552D1^* and *col4a1^G552D2^* lines did not complement each other, but complemented the other variants by interallelic complementation (Kelemen-Valkony et al., 2012).

The series of mutations proved to cover the collagenous region of the *col4a1* gene that corresponds to amino acid 233 up to 1393 within the 170 kDa COL4A1 protein. The *col4a1^G233E^* allele is within the peptide GFP**G/E**EKGERGD (the G to E substitution in bold), a putative integrin binding site in the COL4A1 protein (Parkin et al., 2011). The mutation sites in the *col4a1^G552D1^* and *col4a1^G552D2^* lines localize in the immediate proximity of the peptide GLPGEKGLRGD that resembles the integrin binding site in the COL4A2 protein in the triple helical model made up of (COL4A1)_2_COL4A2 protomers in *Drosophila*, as proposed by us (Kelemen-Valkony et al., 2012).

### Aberrant muscle fiber morphology in *col4a1* mutants

The striated muscles of the common oviduct in all mutants were analyzed by confocal fluorescence microscopy. The most conspicuous phenotype of the mutants was the loss of sarcomeres at restrictive temperature of 29 °C (Fig. 1, B4 through G4, Supplementary Fig. S2, A4, B4), whereas in wild-type control flies normal sarcomere structure and striation was present at both 20°C and 29°C (Fig. 1, A1, A4). In mutants at 29°C, parallel ordered enhanced actin staining intensity areas were present within the muscle fibers that extended over areas larger than a single sarcomere, resembling actin stress fibers or excess actin cross-linking (Fig. 1, white rectangles in B4 through G4, Supplementary Fig. S2, A4, B4). Beyond these areas, amorphous, intensive actin staining aggregates appeared in the sarcoplasm (Fig, 1, white arrows in B4 through G4, Supplementary Fig. S2, A4, B4). An additional prominent phenotype of the *col4a1* mutants at 29°C was the irregular and uneven COL4A1 deposition in the individual muscle fibers (Fig. 1, white arrowheads, B5 through G5, Supplementary Fig. S2, A5, B5), while in wild-type controls homogenous COL4A1 staining was present at 29°C (Fig. 1, A5). In the isoallelic mutants *col4a1^G552D2^* and *col4a1^G1025E2^* the same COL4A1 staining pattern was observed (Supplementary Fig. S2) as in the other lines of the allelic series (Fig. 1), therefore we conclude that the compromised sarcoplasmic morphology and uneven COL4A1 staining/localization is a general phenotype of our series of *col4a1* mutations.

**Fig. 1.**
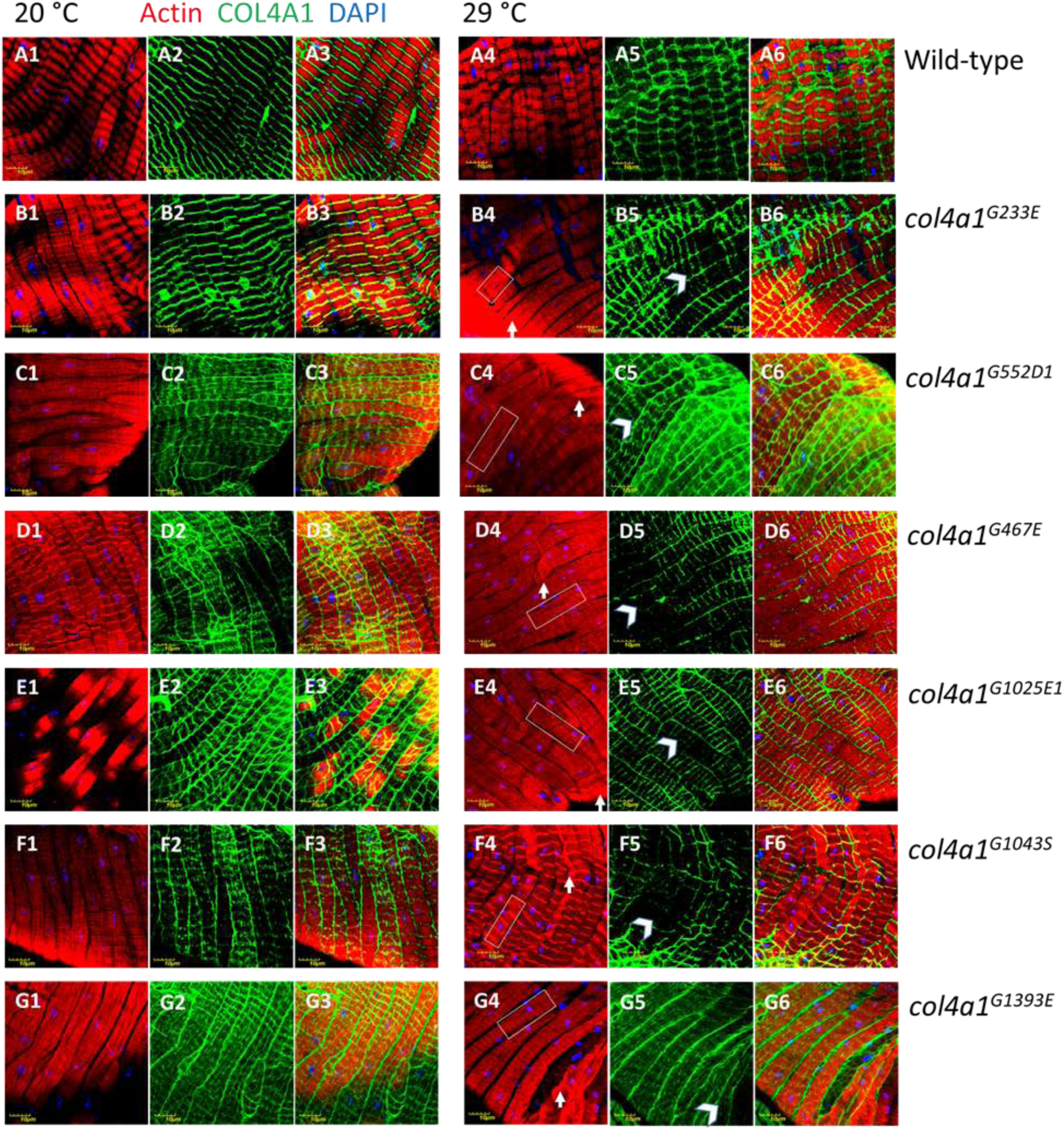
Loss of sarcomeres in *col4a1^G233E^*, *col4a1^G467E^*, *col4a1^G552D1^*, *col4a1^G1025E1^*, *col4a1^G1043S^* and *col4a1^G1393E^* mutant lines at 29 °C (B4 through G4), representative actin bundles (white rectangles in B4 through G4), actin aggragates (white arrows in B4 through G4), uneven COL4A1 expression (white arrowheads in B5 through G5). A3 through G3 and A6 through G6: Overlays of actin and COL4A1 stainings.

### The muscle phenotype co-segregates with mutation-carrying chromosome

The pair of *col4a1* and *col4a2* genes localize to the 25C band of the second chromosome in *Drosophila*. If two different dominant temperature-sensitive (DTS) mutations are present in *trans* configuration the compound heterozygotes provide viability by interallelic complementation (Kelemen-Valkony et al., 2012). In order to determine to what extent compound heterozygotes can recapitulate the dominant temperature-sensitive phenotype affecting sarcoplasmic actin morphology, we have generated the *col4a1^G233E/G1025E1^* double mutant and its reciprocal pair *col4a1^G1025E1/G233E^*. In both compound heterozygotes, the sarcomeric structure was lost, actin bundles developed (Fig. 2, A, D, white rectangles), intensively staining actin aggregates were deposited (Fig. 2, A, D, white arrows), and the COL4A1 protein was detected in uneven and irregular pattern (Fig. 2, B, E, white arrowheads). These results indicate that in *col4a1 ^+/-^* heterozygotes, compromised sarcoplasmic actin morphology and aberrant COL4A1 expression and localization are linked to *col4a1* mutations, are independent from the genetic context, and are not a secondary effect of increased temperature.

**Fig. 2.**
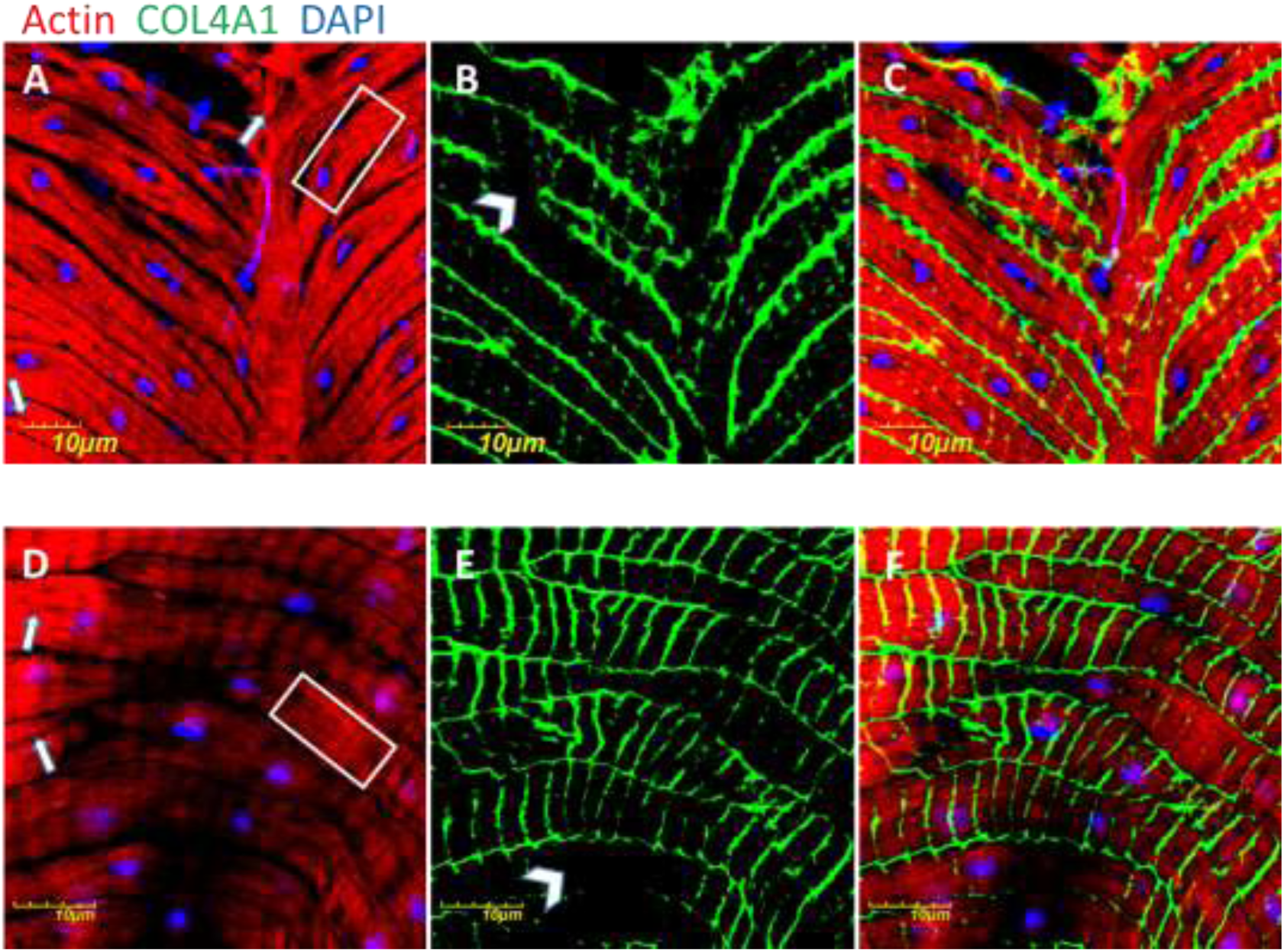
Sarcomeric loss (A D), actin bundles (A, D, white rectangles), intensively staining actin aggregates (A, D, white arrows), COL4A1 protein in uneven and irregularly deposited fashion (B, E, white arrowheads) in compound heterozygotes. C, F: Overlays of actin and COL4A1 stainings.

### Z-disc disintegration, streaming and aberrant integrin expression

In muscles of wild-type *Drosophila*, integrin is expressed at the muscle attachment sites and appears as punctate staining at the costameres aligned with Z-discs (Rui et al., 2010). In order to determine the exact position of the Z-discs in the muscle fibers of the oviduct, we used antibodies against the scaffold protein kettin as a morphological marker. In wild-type controls kettin staining appeared as parallel-ordered lines in each fiber perpendicular to the long axis delineating the sarcomeres and proper striation at both permissive and restrictive temperatures (Fig. 3, A1, A4). Immunohistochemistry using anti-integrin antibodies provided the same staining pattern in close localization as observed for kettin (Fig. 3, A2, A5, and overlays (Fig. 3, A3, A6), confirming integrin localization to the Z-discs in muscle fibers of the oviduct.

**Fig. 3.**
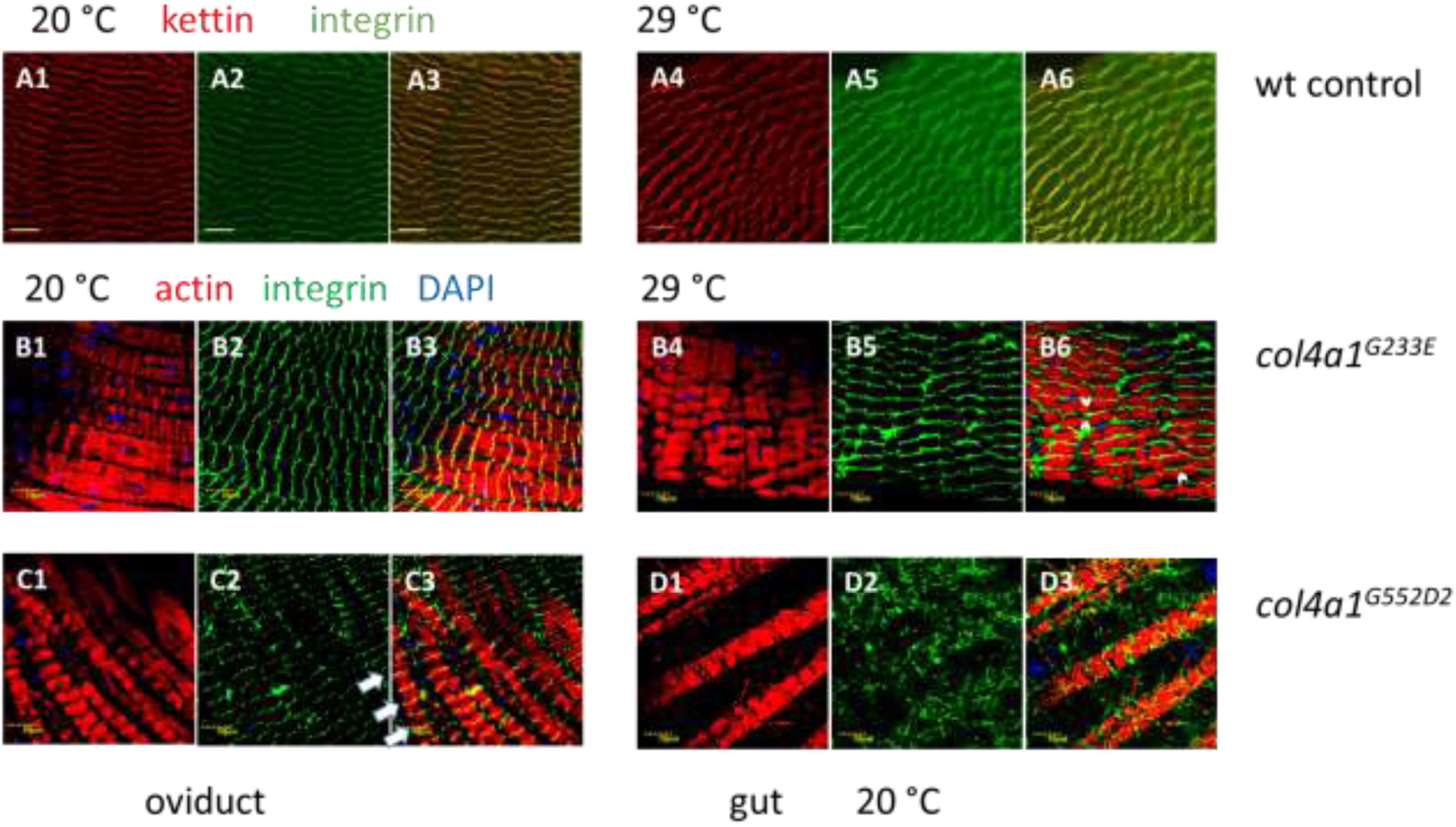
Staining of muscle fibers by the Z-disc marker kettin (A1, A4) and by integrin (A2, A5) and their close localization (overlays A3, A6) in wild-type controls at 20 and 29 °C. Disintegration, streaming of the Z-discs (B5 vs. A2), Z-disc shifting towards the A-band (B6, white arrowheads), integrin deposition at the level of M-disc (C3, white arrows), integrin misexpression in the interfibrillar regions (D3) in the mutants.

In *col4a1* mutant, we observed aberrant integrin expression in the epithelial cells of the Malpighian tubules (Kiss AA et al., 2016), and also surmised irregular integrin deposition in muscle fibers. In mutant oviductal muscle fibers the Z-disc structure, delineated by integrin expression, was disrupted and formed a zig-zag pattern, Z-disc material appeared torn across a large part of sarcomere, and integrin staining was deposited randomly within the sarcomere as dots, consistent with the muscle pathology of Z-disc streaming (Fig. 3, B2, B5). These phenotypic features were enhanced by incubating the mutants at restrictive temperature (Fig. 3, B2 vs. B5). In milder phenotypic manifestation at a low penetrance, ectopic assembly of Z-discs was noted by transition toward the anisotropic (A) band (Fig. 3, B6, white arrowheads) and transition of the Z-disc to the middle of A-band, at the level of M-discs (Fig. 3, white arrows, C3). In these muscle fibers the normal I(Z)-A-I(Z) register of the Z-discs is pushed toward the A(Z)-I-A(Z) pattern (Fig. 3, white arrows, C3). In the muscle of *C. elegans* the M-discs and the dense bodies (Z-disc analogs) are known to function in transmitting the force of muscle contraction to the hypodermis and cuticle, allowing movement of the animal and demonstrating muscle-BM attachment at the level of the M-discs (Qadota and Benian, 2010).

Z-disc streaming, similar to erroneous actin deposition and morphology, seems to be a general feature of *col4a1* mutants, as the same phenotype was observed in each member of the allelic series under restrictive condition (Supplementary Fig. S3).

As *COL4A1* mutations are known to associate with a systemic phenotype affecting multiple tissues and organs (Jeanne and Gould, 2017), we also evaluated muscle fiber morphology in the gut, and at a different developmental stage, in L3 larvae. Integrin deposition was detected in the interfibrillar regions in the mutants (Fig. 3, D2, D3), where no integrin deposition occurred in wild-type L3 larval gut (Kiss M et al., 2016).

### Fiber atrophy and fiber size diversity

The diameter of the individual muscle fibers were measured and the most frequent value was found to correspond to 8 μm both in mutant lines and wild-type controls, incubated at permissive temperature (Fig. 4, Supplementary Fig. S4, Table 2). Incubation of mutant animals at 29 °C shifted the diameters of the muscle fibers toward smaller values. The ratio of the muscle fibers with diameters below 8 μm and down to 4 μm increased by 12-34 %, whereas the same ratio in control flies remained 30% at both temperatures with the majority of fiber diameters in the range of 7-8 μm and only 4-6 % of 6 μm as the smallest value (Fig. 4A, Supplementary Fig. S3, Table 2). We tested muscle fiber features by means of optical sectioning using fluorescent confocal microscopy by gradually lowering the position of the focus. We detected uneven surface and atrophic areas within individual fibers in mutants (Fig. 4B). Collectively, the data showed size heterogeneity, wasting and atrophy of muscle fibers, characteristic features of dystrophic muscle in the *col4a1* mutant lines.

**Fig. 4.**
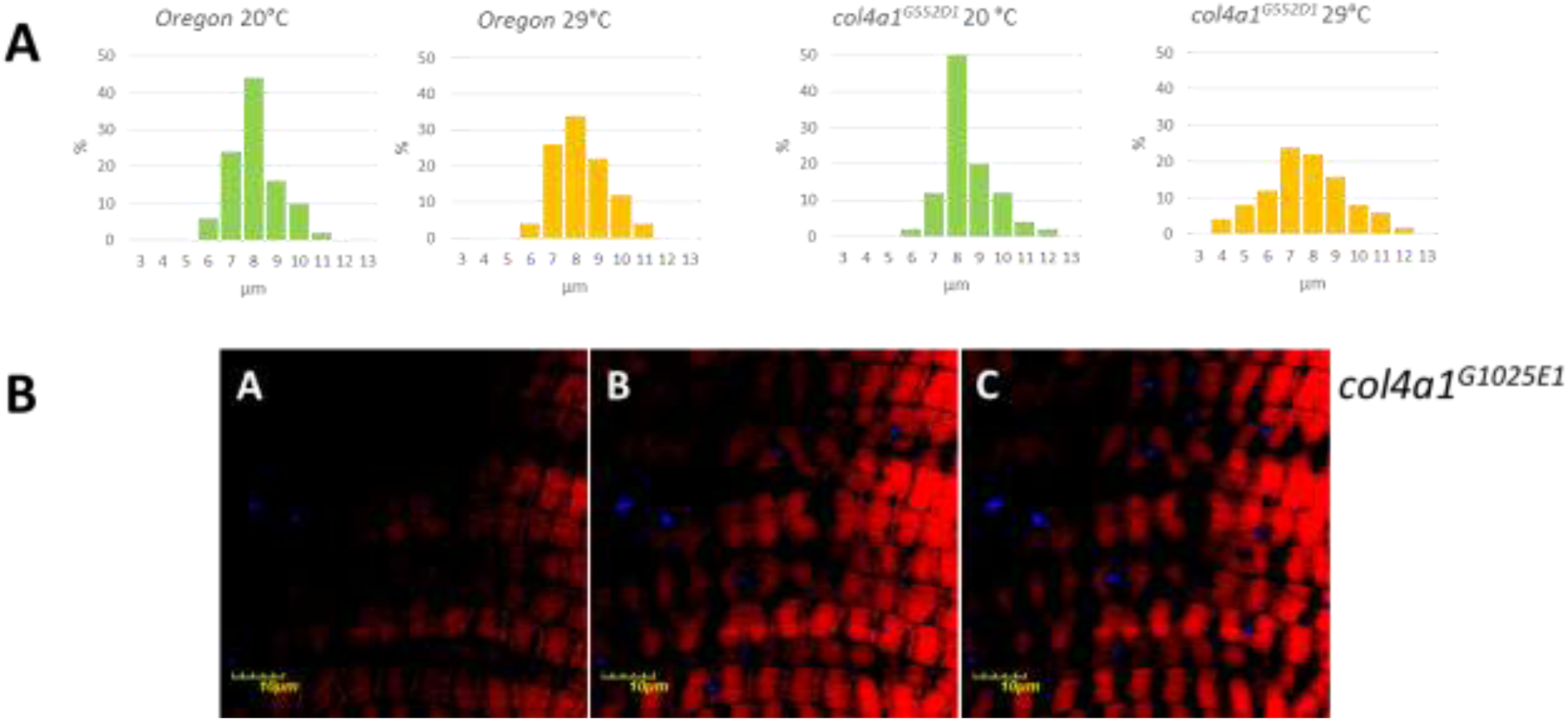
A: Fiber atrophy in the mutant measured by the reduced diameter of the muscle fibers in the mutants at 29 °C. B: Uneven surface and atrophic areas within individual mutant fibers revealed by lowering of the focus of the confocal fluorescence microscope.

**Table 2.**
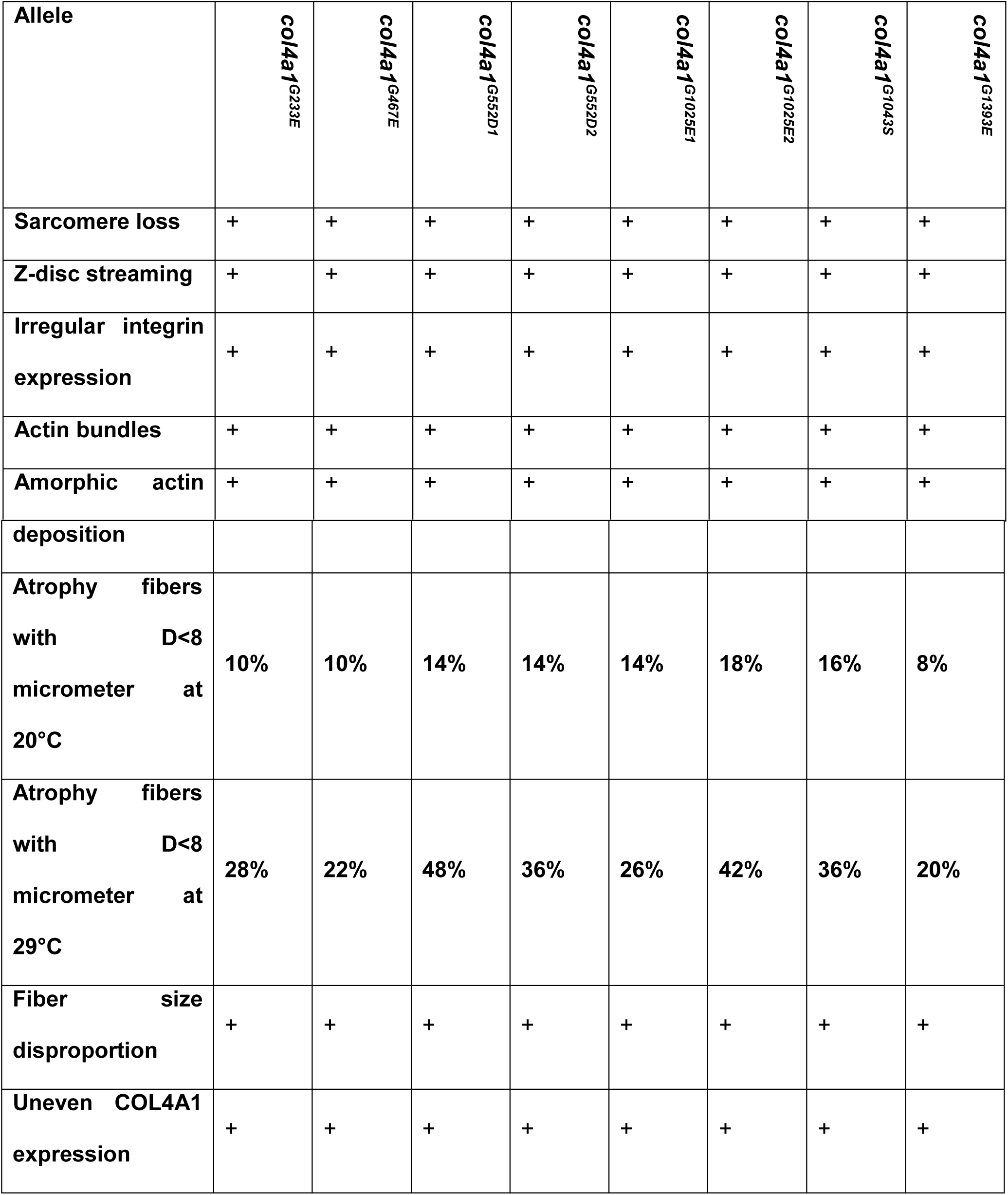
Summary of the phenotypes of the *col4a1* mutants.

## DISCUSSION

In our mutant series, glycine substitutions by large, charged or polar amino acids, glutamate, aspartate and serine, occurred within the *col4a1* gene by transition of the second guanine nucleotide to adenine consistent with the mutagen EMS. Two isoallelic variants *col4a1^G552D1^, col4a1^G552D2^*, and *col4a1^G1025E1^, col4a1^G1025E2^* alleles were identified. The importance of the Gly552 residue is reflected by the fact that a recent EMS mutagenesis resulted in the isolation of the same, temperature-sensitive, dominant-negative *col4a1^G552D^* allele (Hollfelder et al., 2014). Genotype–phenotype relationships explored in over hundred *COL4A1* mutants identified in patients and in murine models revealed that the position of the mutation and not the biochemical properties of the substituting amino acid seems to have a greater impact on the phenotype and disease severity (Jeanne and Gould, 2017). In our *Drosophila* mutant series, however, regardless of the position of the mutation within the collagenous domain of the *col4a1* gene, we observed similar phenotypic defects.

As the oviduct, and also the larval body wall muscle phenotype (Kelemen-Valkony et al., 2012), compromised actin organization and deposition, loss of sarcomere structure were noted in all alleles studied, features that are common in myopathic or dystrophic conditions, with disintegrating muscle sarcomeres together with disintegration and streaming of Z-discs (Rahimov and Kunkel, 2013). These morphologic changes impact the function of the common oviduct as females become sterile and do not lay eggs (Kelemen-Valkony et al., 2012). Conditional knockdown of genes in *Drosophila*, involved genes in integrin mediated adhesion, including *talin*, *alpha-actinin*, *integrin-linked kinase*, *alpha PS2* and *beta PS* integrins result in the common phenotype of Z-disc streaming (Perkins et al., 2010), similar to our *col4a1* mutant series, indicating functional interdependence. Walker-Warburg Syndrome is diagnosed as a monogenic trait in patients carrying mutations in several genes, including *POMT* loci beyond *COL4A1* (Labelle-Dumais et al., 2011). Importantly, in *Drosophila* mutants defective in protein O-mannosyltransferases the symptoms of the Walker-Warburg Syndrome were identified; as part of the mutant phenotypes Z-disc streaming, actin filament disorganization and bundle formation were reported also (Ueyama et al., 2010). The experimental observations clearly indicate the usefulness of the *Drosophila* model in these conditions.

The extensive list of human myopathic/dystrophyc conditions marked by Z-disc streaming and sarcomeric disorganization is missing genes that encode components of integrin-mediated adhesion markedly by genetic reasons. The *Ilk^-/-^* mouse embryos die during periimplantation stage due to impaired epiblast polarization and F-actin accumulation at integrin attachment sites (Sakai et al., 2003). Knockout mutants for the integrin beta subunits and for majority of the alpha subunits have been constructed with phenotypes ranging from a complete block in preimplantation, through developmental defects to perinatal lethality, demonstrating the specificity of each integrin. Muscular dystrophy was observed in patients with *ITGA5* or *ITGA7* mutations (Hynes 2002). Z-disc streaming, however, was not reported in association of *ITGA5* or *ITGA7* mutations. Mouse mutants of *talin 1* or *talin 2* perform myopathy and disassembly of the sarcomeres (Conti et al., 2009). However, the embryonic lethal mutation in the *Drosophila rhea* gene encoding talin recapitulate the phenotype of the integrin beta PS mutations, demonstrating their functional similarities (Brown et al., 2002). These results indicate that genes involved in integrin-mediated adhesion are essential, their homozygous recessive or null mutations are often lethal. The conditional lethality of the temperature-sensitive, heterozygous *col4a1 Drosophila* mutant series allowed manifestation of phenotypic elements that would be non-explorable in humans or mice, such as the disrupted sarcomeric cytoarchitecture and Z-disc streaming that support a role for COL4A1 in integrin mediated adhesion. In conclusion, our *Drosophila* mutant series may serve as an effective model to uncover the mechanisms by which *COL4A1* mutations result in disrupted myofiber-basement membrane interactions and compromised muscle function.

## MATERIALS AND METHODS

### PCR amplification and sequencing

The algorithm of Primerfox was used to design sequence specific primers for the *col4a1* gene. The sequences of the forward and reverse primer pairs are displayed in Table 3. The amplification reaction was carried out with the aid of KAPA Taq polymerase and Fermentas dNTP mix (Thermo Scientific) guided by a touchdown PCR protocol. The initial denaturation at 94 °C lasted for 150 sec, followed by 30 cycles of 93 °C for 15 sec, then 65 °C (−0.6 °C/cycle) for 15 sec and 72 °C for 45 sec. The final elongation step at 72 °C was allowed to run for 180 sec. The lengths of the products were checked on 1% agarose gel, followed by cleanup on silica columns (ZenonBio, Szeged, Hungary). The DNA samples containing the appropriate primers were sent to Eurofins Genomics for sequencing. The same PCR fragments originating from different reactions were read multiple times in both directions to ensure reliability of the results. The received sequence information was aligned to the database of NCBI with the Blast algorithm.

**Table 3.**
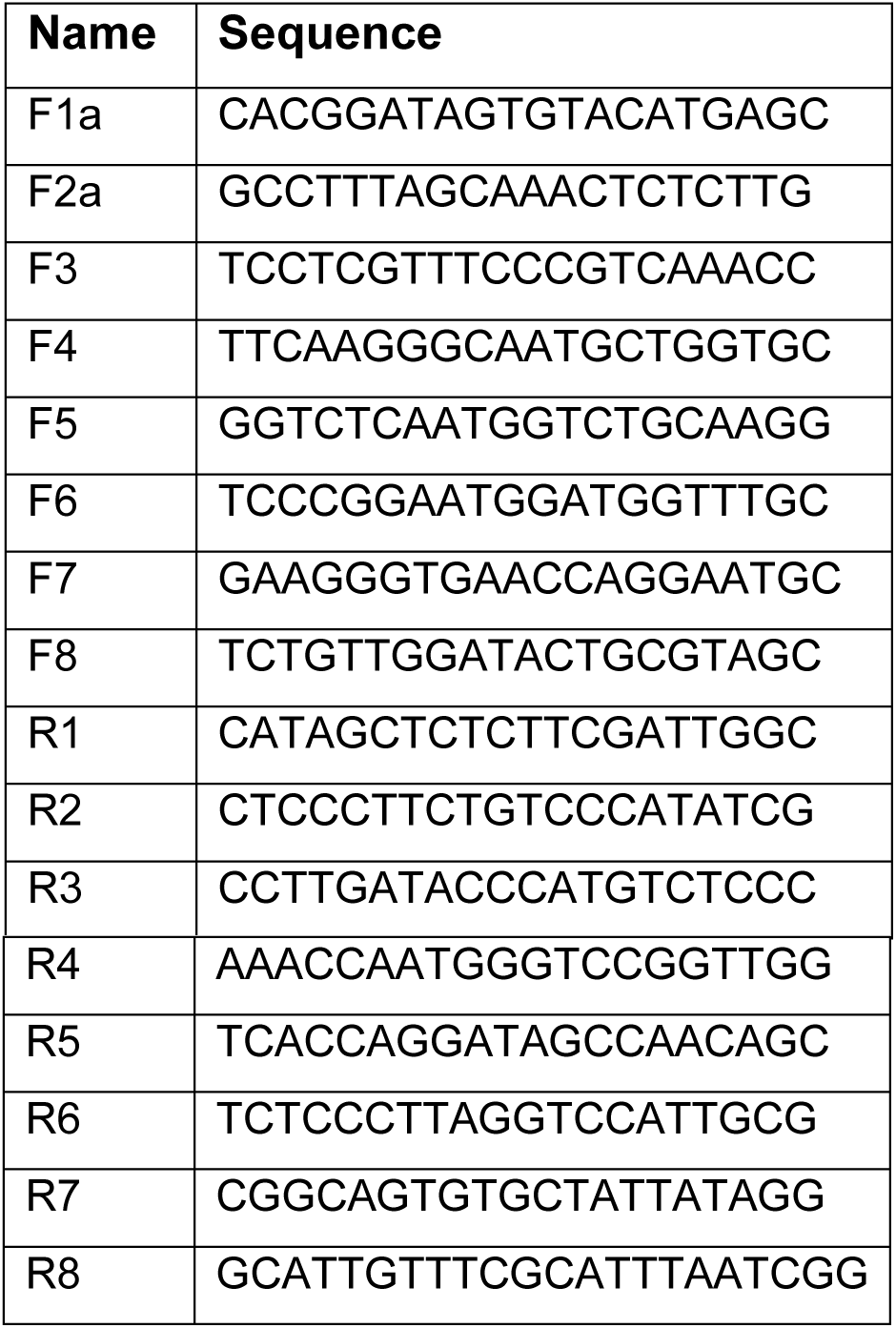
List of forward and reverse PCR primer pairs used for sequencing the *col4a1* gene.

### Maintenance of *Drosophila* strains

Wild-type *Oregon* flies and *col4a1* mutant stocks were maintained at 20°C and 29°C on yeast-cornmeal-sucrose-agar food, completed with the antifungal nipagin. The mutant stocks were kept heterozygous over the *CyRoi* balancer chromosome that prevents recombination with the mutation-carrying homolog. Common oviducts were removed under carbon dioxide anesthesia from adults that were grown at both permissive and restrictive temperature for 14 days. The common oviduct is circumfered by a single-layer of striated muscle fibers thus sectioning can be avoided and sarcolemma-associated events can directly be observed. Dissected common oviducts were fixed in 4% paraformaldehyde dissolved in phosphate buffered saline (PBS) for 10 min, washed tree times in PBS, permeabilized for 5 min in 0,1% Triton X dissolved in PBS and washed tree times in PBS. Blocking was achieved for in 5% BSA dissolved in PBS for 1 hour, and washed tree times in PBS. Trans-heterozygous strains were generated by crossing two *col4a1^+/-^* heterozygotes selecting for the loss of the balancer chromosome *CyRoi*.

### Immunostaining and antibodies

Nuclei in the dissected common oviducts were counter-stained by 1μg/ml 4’,6-diamino-2-phenylindol (DAPI) in 20 μl PBS, 12 min in dark. F-actin was stained by 1 unit Texas Red^TM^-X Phalloidin (ThermoFisher) in 20 μl PBS for 20 min. Integrin dimer staining was achieved by an equimolar mixture consisting of both anti-integrin monoclonal antibodies (mouse, Developmental Studies Hybridoma Bank) that recognize alpha PS I or alpha PS II subunits. Mouse antibody against *Drosophila* COL4A1 protein was generated by Creative Ltd, Szeged, Hungary. Primary mouse antibodies were visualized by 1 μl F(ab’) 2-Goat Anti-Mouse IgG (H+L) Cross Adsorbed Secondary Antibody conjugated with Alexa Fluor 488 (ThermoFisher) in 20 μl PBS for 1 hour or 1 μl Goat Anti-Mouse IgG (H+L) Cross Adsorbed Secondary Antibody, Alexa Fluor 350, in 20 μl PBS for 1 hour.

### Confocal microscopy

Photomicrographs of the common oviducts were generated by confocal laser scanning fluorescence microscopy (Olympus Life Science Europa GmbH, Hamburg, Germany). Microscope configurations were set up as described (Kiss AA et al., 2018). Briefly, objective lens: UPLSAPO 60x (oil, NA: 1.35); sampling speed: 8 μs/pixel; line averaging: 2x; scanning mode: sequential unidirectional; excitation: 405 nm (DAPI), 543 nm (Texas Red) and 488 nm (Alexa Fluor 488); laser transmissivity: 7% were used for DAPI, 42% for Alexa Fluor 488 and 52% for Texas Red.

### Size determination of the muscle fibers

Five confocal photomicrographs displaying oviducts were stained by Texas Red^TM^-X Phalloidin and anti-COL4A1 antibody, taken from all mutants and wild-type controls at 20 and 29 °C. Diameters of ten randomly chosen muscle fibers were measured generating altogether 900 values. Bins of diameter intervals differing by one μm were displayed in histograms showing the numbers of the corresponding diameters.

## Author Contributions

Conceived and designed the experiments: M.M. Performed the experiments: A.A.K., NP, M.K., M.M. Analyzed the data: A.A.K., N.P., M.K., M.M. Provided resources: ZB. Writing - original draft: M.M., C.K.; Writing - review & editing: M.M., C.K., Z.B.; Supervision: M.M.; Project administration: M.M., A.A.K.; Funding acquisition: M.M., A.A.K.

---

**Supplementary Fig. S1.**
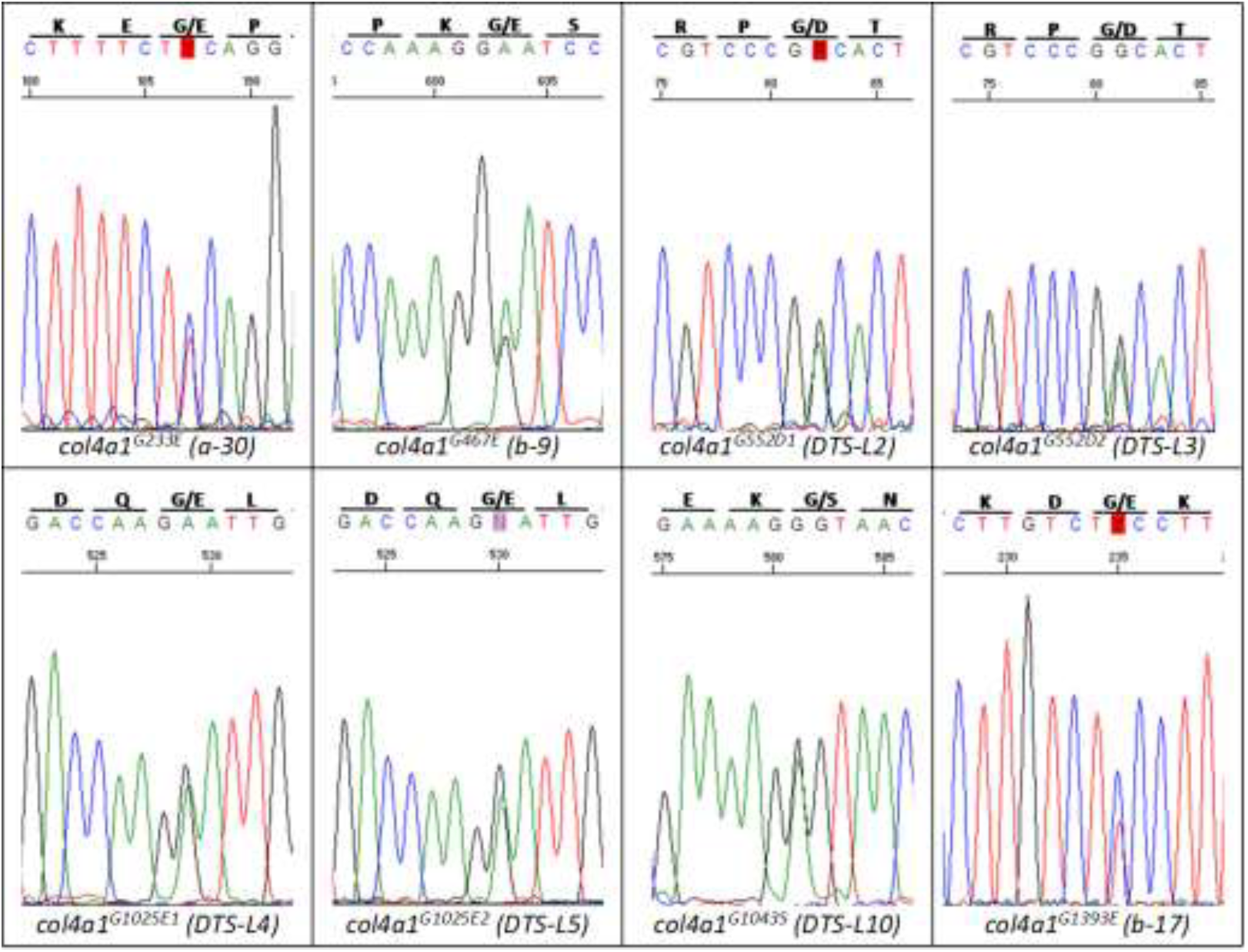
Nucleotide sequence of the heterozygous lesions in *col4a1* alleles. Note the isoallelic mutations G552D and G1025E. For a better resolution the sequence of the complementary strand is displayed in the *b-17* mutant.

**Supplementary Fig. S2.**
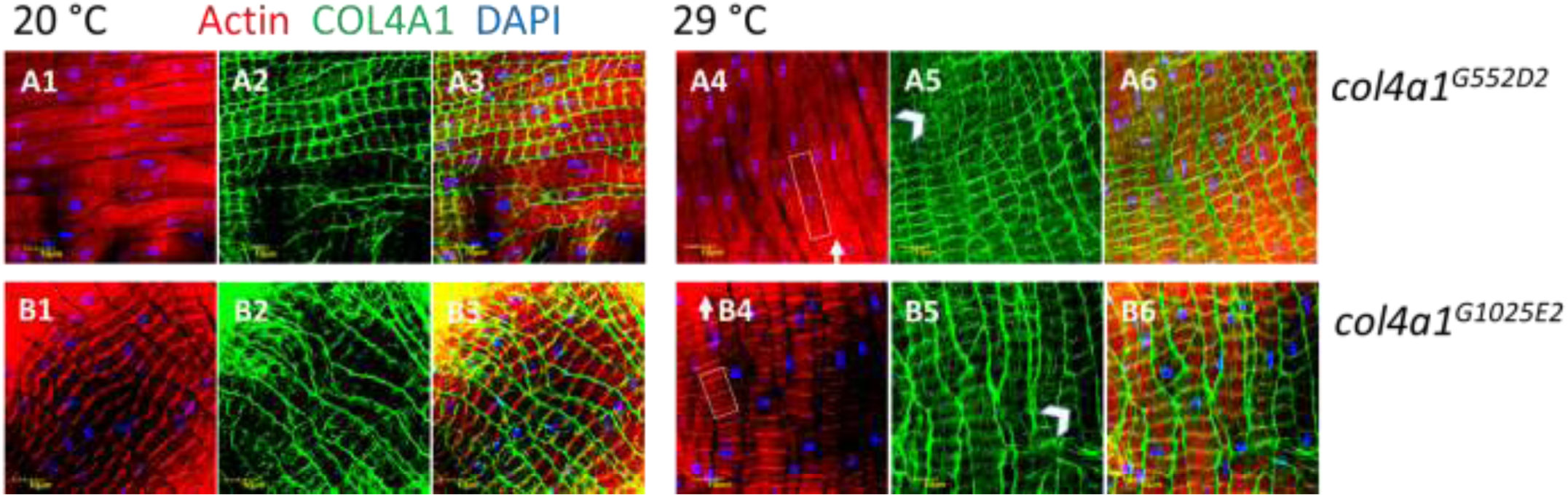
Loss of sarcomeres (A4, B4) actin bundle development (white rectangles, A4, B4), actin aggregates (white arrows, A4, B4), irregular COL4A1 deposition (white arrowheads, A5, B5) in the isoallelic lines *col4a1^G552D2^* and *col4a1^G1025E2^*. A3, B3, A6, B6: Overlays.

**Supplementary Fig. S3.**
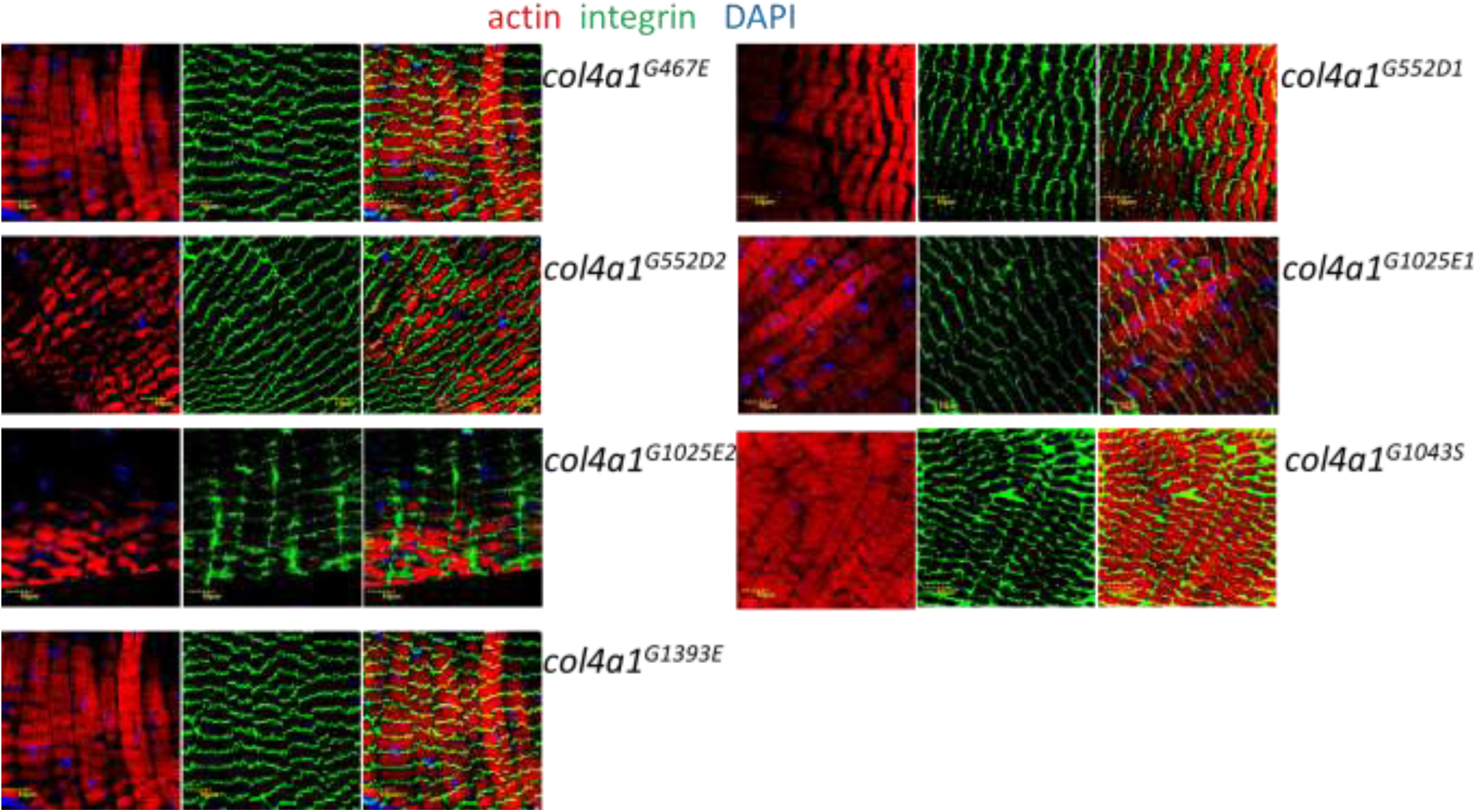
Integrin staining in the *col4a1* mutants at 29 °C. Note uneven deposition, disintegration, zig-zag pattern of integrin expression demonstrating streaming of the Z-discs.

**Supplementary Fig. S4.**
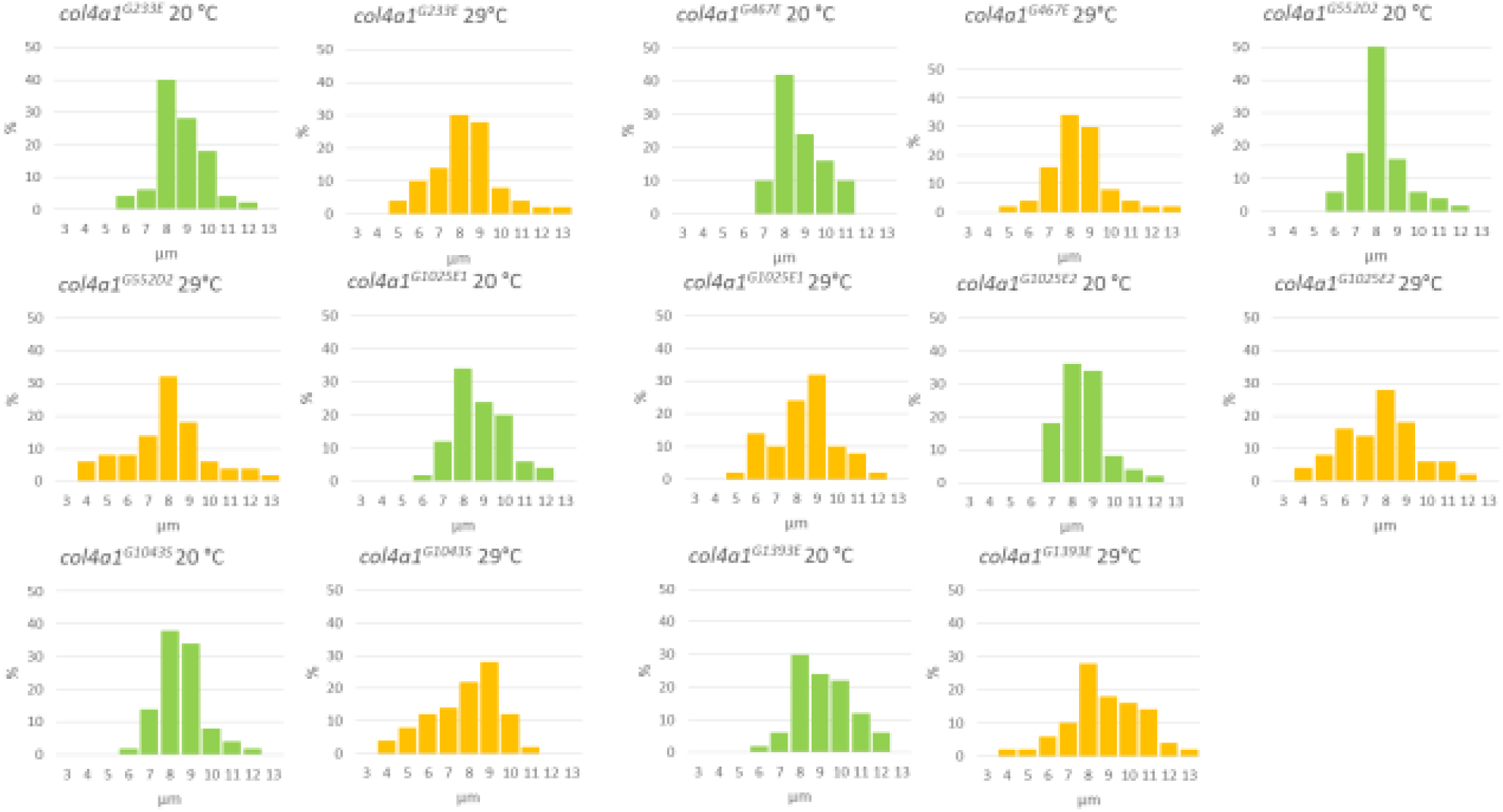
Diameters of muscle fibers are shifted toward reduced values (yellow columns) in mutants at 29 °C.

## References

van Agtmael, T., Schlötzer-Schrehardt, U., McKie, L., Brownstein, D. G., Lee, A. W., Cross, S. H., Sado, Y., Mullins, J. J., Pöschl, E. and Jackson, I. J. (2005) Dominant Mutations of Col4a1 Result in Basement Membrane Defects Which Lead to Anterior Segment Dysgenesis and Glomerulopathy. Hum Mol Genet. 14, 3161–3168.

Alamowitch, S., Plaisier, E., Favrole, P., Prost, C., Chen, Z., Van Agtmael, T., Marro, B. and Ronco, P. (2009) Cerebrovascular Disease Related to COL4A1 Mutations in HANAC Syndrome. Neurology 73, 1873–1882.

Alport, A. C. (1927) Hereditary familial congenital haemorrhagic nephritis. Br Med J. 1, 504–506.

Bertini, E., D’Amico, A., Gualandi, F., Petrini, S. (2011) Congenital Muscular Dystrophies: A Brief Review. Semin Pediatr Neurol. 18, 277–288.

Brown, N. H., Gregory, S. L., Rickoll, W. L., Fessler, L. I., Prout, M., White, R. A. H., Fristrom, J. W. (2002) Talin Is Essential for Integrin Function in Drosophila. Dev Cell 3, 569–579.

Conti, F. J., Monkley, S. J., Wood, M. R., Critchley, D. R., Müller. U. (2009) Talin 1 and 2 are required for myoblast fusion, sarcomere assembly and the maintenance of myotendinous junctions. Development. 136, 3597–3606.

Durbeej, M., Campbell, K. P. (2002) Muscular dystrophies involving the dystrophin–glycoprotein complex: an overview of current mouse models. Curr Opin Genet Dev. 12, 349–361.

Favor J, Gloeckner CJ, Janik D, Klempt M, Neuhäuser-Klaus A, Pretsch W, Schmahl W and Quintanilla-Fend L. (2007) Type IV Procollagen Missense Mutations Associated with Defects of the Eye, Vascular Stability, the Brain, Kidney Function and Embryonic or Postnatal Viability in the Mouse, Mus Musculus: An Extension of the Col4a1 Allelic Series and the Identification of the First Two Col4a2 Mutant Alleles. Genetics 175, 725–736.

Gould, D. B., Phalan, F. C., Breedveld, G. J., van Mil, S. E., Smith, R. S., Schimenti, J. C., Aguglia, U., van der Knaap, M. S., Heutink, P. and John. S. W. M. (2005) Mutations in Col4a1 Cause Perinatal Cerebral Hemorrhage and Porencephaly. Science 308, 1167–1171.

Guicheney, P., Vignier, N., Helbling-Leclerc, A., Nissinen, M., Zhang, X. (1997) Genetics of laminin alpha 2 chain (or merosin) deficient congenital muscular dystrophy: From identification of mutations to prenatal diagnosis. Neuromuscul Disord. 7, 180–186.

Guiraud, S., Migeon, T., Ferry, A., Chen, Zh., Ouchelouche, S., Verpont, M-C., Sado, Y., Allamand, V., Ronco, P., Plaisier, E. (2017) HANAC Col4a1 Mutation in Mice Leads to Skeletal Muscle Alterations due to a Primary Vascular Defect. Am J Pathol. 187(3), 505–516.

Hollfelder, D., Frasch, M., Reim, I. (2014) Distinct functions of the laminin β LN domain and collagen IV during cardiac extracellular matrix formation and stabilization of alary muscle attachments revealed by EMS mutagenesis in Drosophila. BMC Dev Biol. 14, 26.

Hudson, B. G., Tryggvason, K., Sundaramoorthy, M., Neilson, E. G. (2003) Alport’s Syndrome, Goodpasture’s Syndrome, and Type IV Collagen. N Engl J Med. 348, 2543–2556.

Hynes, RO. (2002) Integrins: Bidirectional Allosteric Signaling Machines. Cell, 110, 673–687.

Jeanne, M., Gould, D. B. (2017) Genotype-phenotype correlations in pathology caused by collagen type IV alpha 1 and 2 mutations. Matrix Biol. 57–58, 29–44.

Kelemen-Valkony, I., Kiss, M., Csiha, J., Kiss, A., Bircher, U., Szidonya, J., Maróy, P., Juhász, G., Komonyi, O., Csiszár, K. and Mink, M. (2012) Drosophila basement membrane collagen col4a1 mutations cause severe myopathy. Matrix Biol. 31(1), 29–37.

Kelemen-Valkony, I., Kiss, M., Csiszár, K., Mink, M. (2012A) Inherited Myopathies. In: Myopathies: New Research. Eds: Howard S. Washington and Chris E. Castillo Jimenez, Nova Publishers, ISBN: 978-1-62257-372-1.

Kiss, A. A., Popovics, N., Boldogkői, Z., Csiszár, K., Mink, M. (2018) 4-Hydroxy-2-nonenal Alkylated and Peroxynitrite Nitrated Proteins Localize to the Fused Mitochondria in Malpighian Epithelial Cells of Type IV Collagen Drosophila Mutants. Biomed Res Int. 3502401.

Kiss, A. A., Popovics, N., Szabó, G., Csiszár, K., Mink, M. (2016) Altered stress fibers and integrin expression in the Malpighian epithelium of Drosophila type IV collagen mutants. Data in Brief 7, 868–872.

Kiss, M., Kelemen-Valkony, I., Kiss, A. A., Kiss, B., Csiszár, K., Mink, M. (2012) Muscle dystrophy is triggered by type IV collagen alleles affecting integrin binding sites directly or indirectly in Drosophila. Acta Biochim Pol. 59, 26.

Kiss, M., Kiss, A. A., Radics, M., Popovics, N., Hermesz, E., Csiszár, K. and Mink M. (2016) Drosophila type IV collagen mutation associates with immune system activation and intestinal dysfunction. Matrix Biol. 49, 120–131.

Kuo, D. S., Labelle-Dumais, C., Gould, D. B. (2012) COL4A1 and COL4A2 Mutations and Disease: Insights into Pathogenic Mechanisms and Potential Therapeutic Targets. Hum Mol Genet. 21(R1), R97–110.

Labelle-Dumais, C., Dilworth, D. J., Harrington, E. P., De Leau, M., Lyons, D., Kabaeva, Z., Manzini, M. C., Dobyns, W. B., Walsh, C. A., Michele, D. E., Gould, D. B. (2011) COL4A1 Mutations Cause Ocular Dysgenesis, Neuronal Localization Defects, and Myopathy in Mice and Walker-Warburg Syndrome in Humans. PLoS Genet. 7, e1002062.

Lampe, A.K., Bushby, K. M. (2005) Collagen VI related muscle disorders. J Med Genet. 42, 673–685.

Mayer, U., Saher, G., Fassler, R., Bornemann, A., Echtermeyer, F., von der Mark, H., Miosge, N., Poschl, E., von der Mark, K. (1997) Absence of integrin alpha-7 causes a novel form of muscular dystrophy. Nature Genet. 17, 318–323.

Parkin, J. D., San Antonio, J. D., Pedchenko, V., Hudson, B., Jensen ST and Savige J. (2011) Mapping Structural Landmarks, Ligand Binding Sites, and Missense Mutations to the Collagen IV Heterotrimers Predicts Major Functional Domains, Novel Interactions, and Variation in Phenotypes in Inherited Diseases Affecting Basement Membranes. Hum Mutat. 32, 127–143.

Perkins, A. D., Ellis, S. J., Asghari, P., Shamsian, A., Moore, E. D., Tanentzapf, G. (2010) Integrin mediated adhesion maintains sarcomeric integrity. Dev Biol. 338, 15–27.

Plaisier, E., Gribouval, O., Alamowitch, S., Mougenot, B., Prost, C., Verpont. M. C., Marro, B., Desmettre, T., Cohen, S. Y., Roullet, E., Dracon, M., Michel Fardeau, M. et al. (2007) COL4A1 Mutations and Hereditary Angiopathy, Nephropathy, Aneurysms, and Muscle Cramps. N Engl J Med., 357, 2687–2695.

Plaisier, E., Chen, Z., Gekeler, F., Benhassine, S., Dahan, K., Marro, B., Alamowitch, S., Paques, M. and Ronco, P. (2010) Novel COL4A1 Mutations Associated with HANAC Syndrome: A Role for the Triple Helical CB3(IV) Domain. Am J Med Genet. A, 152A, 2550–2555.

Pozzi, A., Yurchenco, P. D., Iozzo, R. V. (2017) The nature and biology of basement membranes. Matrix Biol. 57–58, 1–11.

Qadota, H., Benian, G. M. (2010) Molecular structure of sarcomere-to-membrane attachment at M-Lines in C. elegans muscle. J Biomed Biotechnol. 2010:864749, 1–9.

Rahimov, F., Kunkel, L. M. (2013) Cellular and molecular mechanisms underlying muscular dystrophy. J Cell Biol. 201, 499–510.

Rui, Y., Bai, J., Perrimon, N. (2010) Sarcomere Formation Occurs by the Assembly of Multiple Latent Protein Complexes. PLoS Genet. 6(11), e1001208. doi:10.1371/journal.pgen.1001208.

Sakai, T., Li, S., Docheva, D., Grashoff, C., Sakai, K., Kostka, G., Braun, A., Pfeifer, A., Yurchenco, P. D. and Fässler, R. (2003) Integrin-linked kinase (ILK) is required for polarizing the epiblast, cell adhesion, and controlling actin accumulation. Genes Dev. 17, 926–940.

Sanes, J. R. (2003) The Basement Membrane/Basal Lamina of Skeletal Muscle. J Biol Chem. 278, 12601 – 12604.

Ueyama, M., Akimoto, Y., Ichimiya, T., Ueda, R., Kawakami, H., Aigaki, T., Nishihara, S. (2010) Increased Apoptosis of Myoblasts in Drosophila Model for the Walker-Warburg Syndrome. PLoS One. 5(7), e11557. doi:10.1371/journal.pone.0011557.

Volk, T., Fessler, L. I., Fessler, J. H. (1990) A role for integrin in the formation of sarcomeric cytoarchitecture. Cell 63, 525–536.

Zhou, J., Mochizuki, T., Smeets, H., Antignac, C., Laurila, P., De Paepe, A., Tryggvason, K. and Reeders. S. T. (1993) Deletion of the Paired Alpha 5(IV) and Alpha 6(IV) Collagen Genes in Inherited Visceral Muscle Tumors. Science 261, 1167–1169.

